## Supplementary Figures for "ZO-1 regulates Hippo-independent YAP activity and cell proliferation via a GEF-H1- and TBK1-regulated mechanosensitive signalling network"

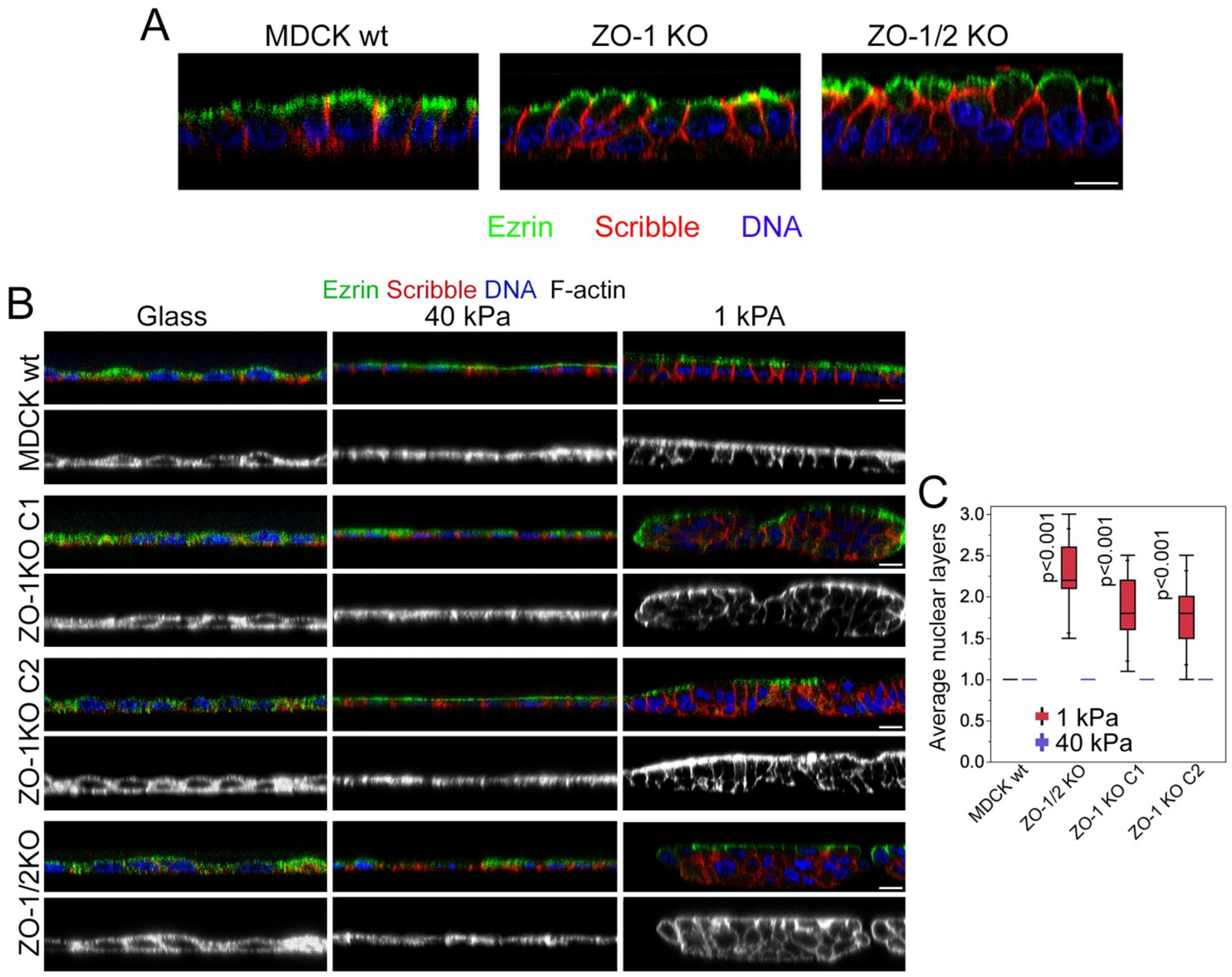


**Supplementary Figure 1. Knockout of ZO-1 increases cell proliferation and disrupts monolayer organization.**

Expression of polarity markers and monolayer organization by control and knockout MDCK cells grown on filters (**A**) or hydrogels (**B**,**C**). Quantification shows means, interquartile ranges, and p-values derived from a signed-rank test (n=15 images per cell line and condition). Magnification bars, 20 μm.

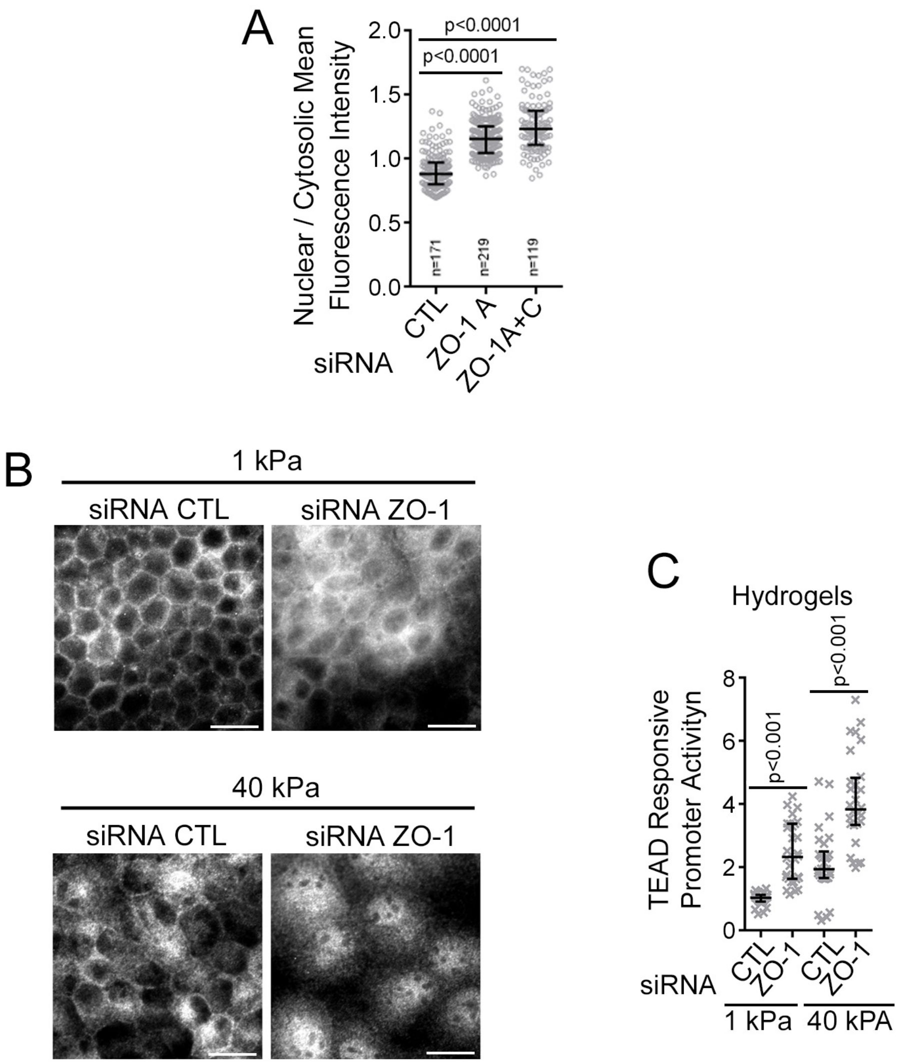


**Supplementary Figure 2. Knockdown of ZO-1 promotes nuclear translocation of YAP.**

**A** Ratio of nuclear to cytosolic YAP was measured in individual cells treated with ZO-1-specific siRNAs (see Fig2F for images). **B** siRNA-transfected MDCK cells grown on 40 and 1 kPa PAA hydrogels were fixed and immunostained for YAP. **C** TEAD transcriptional activity was analyzed by reporter gene assay in MDCK cells grown on hydrogels of 40 and 1 kPa. Quantifications show individual cells analyzed, medians, interquartile ranges, and p-values derived from Wilcoxon tests. Magnification bars, 20 μm.

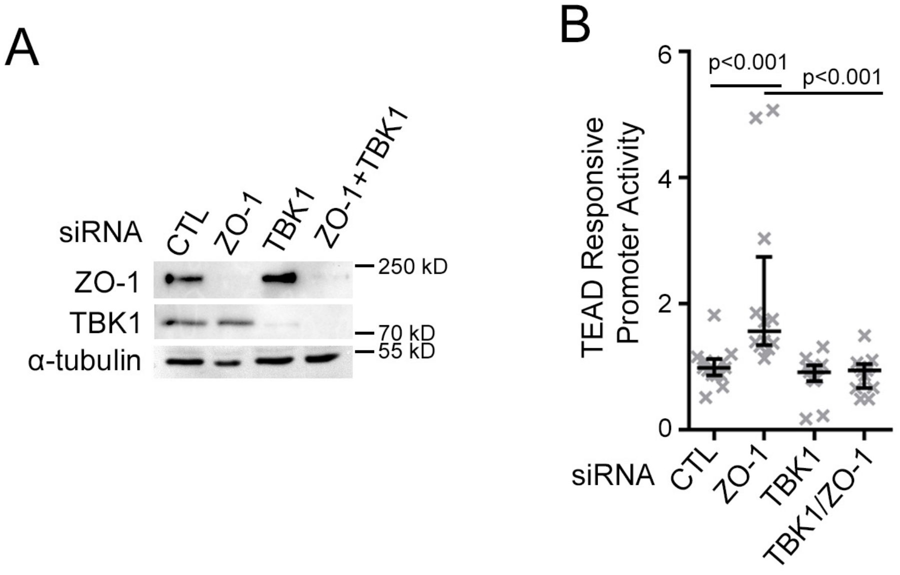


**Supplementary Figure 3. Knockdown of TBK1 inhibits YAP/TEAD activation in ZO-1 depleted cells.**

MDCK cells were transfected with siRNAs as indicated and then analyzed by immunoblotting (**A**) or a TEAD responsive promoter assay (**B**). The reporter assay shows individual determinations, medians, interquartile ranges, and p-values for the indicated pairs from a Wilcoxon assay.

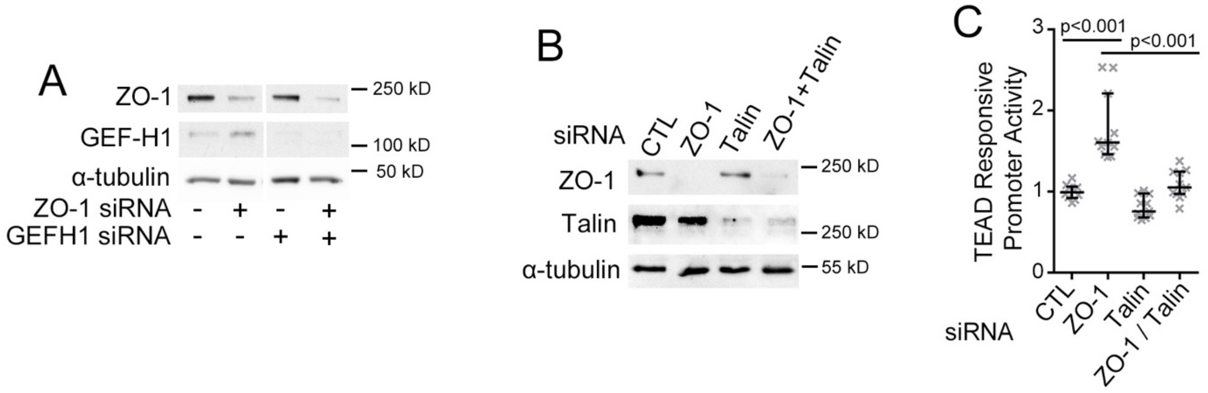


**Supplementary Figure 4. Analysis of siRNA transfected MDCK cells.**

MDCK cells were transfected with siRNAs as indicated and then analyzed by immunoblotting to determine GEF-H1 and ZO-1 expression (**A**), knockdown of ZO-1 and talin (**B**), or the impact on the TEAD responsive promoter gene assay (**C**, shown are individual determinations, medians, interquartile ranges and p-values derived from Wilcoxon tests).
